# Life cycle exposure to cyhalofop-butyl induced reproductive toxicity toward zebrafish

**DOI:** 10.1101/2022.06.14.496026

**Authors:** Tao Zhu, Siwen Wang, Dong Li

## Abstract

Cyhalofop-butyl (CyB) is an herbicide widely used in paddy fields, which may transfer to aquatic ecosystems and cause harm to aquatic organisms. In this study, zebrafish (Danio rerio) were exposed to CyB (0.1, 1 and 10 ug/L) related to the environment throughout its adult life cycle from embryo to sexual maturity. The effects of CyB on zebrafish growth, reproduction and offspring development were studied. It was found that female spawning was inhibited and adult male fertility decreased. In addition, we detected the expression of sex steroid hormones and genes related to hypothalamus-pituitary-gonad-liver (HPGL) axis. After 150days of exposure, the hormone balance of parent zebrafish (F0) was disturbed and the concentrations of 17β-estradiol (E2) and vitellogenin (VTG) in zebrafish were decreased. F1 embryos showed abnormal developmental results, including decreased heart rate, decreased body length, spontaneous motor inhibition, while the developmental abnormalities of F1 embryos were relieved when exposed to CyB-free clear water. The change of sex hormone is regulated by gene expression related to HPGL axis. These results confirmed that long-term exposure to CyB in the environmental concentrations can damage the reproductive capacity of F0 generation zebrafish by disrupting the transcription of genes related to HPGL axis, which may lead to abnormal development of F1. Overall, these data may provide a new understanding of the reproductive toxicity of zebrafish parents and offspring after long-term exposure to CyB.

**Highlights:**

- Environmental level of CyB exposure caused gonadal impairment.
- CyB exposure suppressed spawning ability of zebrafish.
- CyB exposure changed the plasma hormone level of zebrafish and altered HPGL axis in both genders.
- Parental CyB exposure led to abnormal development of F1.

## 1. Introduction

Cyhalofop-butyl (CyB) was first developed by Dow AgroSciences in 1987, which is an aryloxyphenoxypropionate (APP) herbicide widely used in paddy fields all over the world (Bleau et al., 1996; Cao et al., 2016a). It was first introduced to Asia in 1995, is widely sold in rice growing areas (Sondhia and Khare, 2014; Wu et al., 2014) and has been used in agriculture for more than twenty years (Kalsing et al., 2017). In China, it has been registered as an herbicide for selectively controlling weeds in rice fields since 2006, and is increasingly used in agricultural production (li et al., 2020; Shang et al., 2019; Yuan et al., 2019), which leads to its easy transfer to the aquatic environment. Therefore, the potential hazards of CyB that may spread to aquatic organisms should be to taken into account. The detection concentration of CyB in bulk drainage water samples in Japan is 0.01 ∼ 0.08 μg/L (Cao et al., 2016a; Phong et al., 2010). The situation is even worse in China, after spraying 135 g a.i./ha of 10 % fenoxaprop-pethyl•cyhalofop-butyl EC herbicide, the concentration of CyB can be detected as 2.017mg/L in rice fields in southern China (Guo et al., 2008). Some studies have reported the effects of CyB on aquatic organisms. For example, CyB can induce developmental toxicity, oxidative stress and apoptosis of zebrafish embryos (Cao et al., 2016a; Cheng et al., 2021; Zhu et al., 2015), as well as other aquatic organisms, such as *Yellow River Carp* (Xia et al., 2018), *Rana chensinensis tadpole* (Wu et al., 2011) and *Misgurnus anguillicaudatus* (Shang et al., 2019). Therefore, the reproductive toxicity of CyB to aquatic organisms, especially fish, and its potential mechanism are very limited.

Measurement of steroid hormones in teleost is very useful for monitoring the reproductive system because these hormone changes play a vital role in fish reproduction, and correspond to genes involved in the steroid production pathway (Devlin and Nagahama, 2002; Ji et al., 2013b). Moreover, reproductive capacity and individual development are coordinated by the interaction of genes on the hypothalamus-pituitary-gonad-liver (HPGL) axis (Hilscherova et al., 2004; Plant, 2015; Villeneuve et al., 2008; Xu et al., 2011). Changes in sex steroid concentrations may cause reproductive dysfunction such as sex differentiation, fertility rate and fertilization rate decline (Liu et al., 2018). Gonadotropins secreted by the pituitary gland (such as follicle-stimulating hormone, FSH; Luteinizing hormone, LH) act on gonads to regulate sex hormones synthesis and gametogenesis to guide steroid production and gametogenesis. In organisms, FSH and LH regulate the synthesis of 17β-estradiol (E2) and testosterone (T), which participate in gametogenesis and oviposition (Kaiser and Ursula, 2011; Son et al., 2022). In zebrafish gonads, *cyp17* converts 17-hydroxyl progesterone into androstenedione catalyze, which then catalyzed by *17βhsd* to form T (Fernandes et al., 2011). T secreted by follicle is converted into estradiol by aromatase (cyp19a) (Clelland and Peng, 2009; Trant et al., 2001). Therefore, E2 and T disorders will affect fish reproduction. E2 enters the liver through the blood circulation system, stimulating and promoting the synthesis and secretion of vitellogenin (VTG) (Busby et al., 2010). VTG travels through the bloodstream to the ovary, where it is decomposed into yolk protein, which provides sufficient nutrients for the growth and development of offspring. The synthesis and secretion of these steroid hormones promote the sex differentiation of organisms, the growth and development of gonads, and finally regulate their reproductive system (Nagahama and Yamashita, 2008; Sofikitis et al., 2008).

Our research aims at evaluating the reproductive toxicity of CyB to zebrafish, exposing zebrafish embryos (2 hours post fertilization) to the nominal concentrations (0, 0. 1, 1 and 10 μg/L) of CyB for 120 days until adulthood, and revealing the potential impact of CyB on the whole life cycle of zebrafish. The reproductive capacity, sex gland index (GSI), sex steroid hormone and plasma VTG concentrations, gonad histology and relative mRNA levels of genes related to HPGL axis were measured. In addition, the differences of female and male zebrafish responses to CyB were discussed, and the effects of CyB on the growth and development of offspring were studied. The results of this study contribute to better understanding of the negative effects of CyB on fish reproduction and its underlying mechanisms.

## 2. Materials and methods

### 2.1. Chemicals

Cyhalofop-butyl (CAS: 122008-85-9) (97.5%) was obtained from the Jiangsu Zhongqi Technology Co., Ltd (Jiangsu, China) and dissolved in dimethyl sulfoxide (DMSO) as a stock solution. All other reagents were analytical grade.

### 2.2. Zebrafish rearing and CyB exposure

The parental zebrafish of wild-type AB strain were purchased from Hebei Aquarium Department, China. After domestication for two weeks, the parents were cultured in flow-through feeding facility, and at a temperature of 27±1 □, photoperiod of 14:10 h (light: dark), feed with live brine shrimp three times a day. Embryo exposure experiments were performed according to OECD guidelines and methods described in previous studies (Duan et al., 2021; OECD, 1998), and ensure the final DMSO volume is less than 0.01% (v/v). Normal embryos (F0) obtained by mating parent zebrafish in spawning box were randomly assigned to four treatment groups (0, 0. 1, 1 and 10 μg/L) and exposed for 120 days, and changed the exposure liquid every day. After 120 days of exposure, 16 pairs of zebrafish in each treatment group were randomly selected for reproduction test, and four replicates were set at the same time. Eggs are laid once a week, and embryos are collected and counted one hour after fertilization. In addition, the F0 generation oviposited embryos (F1 generation) were collected, and the development status was monitored in standard water until 96 hpf. All animal experiments were conducted under the policy of Animal Ethics Committee of Hebei University.

At the end of spawning experiment (150 d), the spawning fish were fasted for 24 hours and euthanized with 0.03% MS-222. After, collecting the blood of female and male zebrafish, and recording body weight, body length, and the weights of brain, liver, and gonad tissues, samples were frozen at -80□ (each group had three replicates n=5 per replicate) for subsequent gonad histopathological examination and RNA expression analysis, as well as brain body index (BSI), liver body index (HSI) and gonad index (HSI). The collected blood samples will be used to determine sex steroid hormones (E2 and T) and vitellogenin (VTG) levels.

### 2.3 F1 offspring embryo development and water recovery test

The collected F1 embryos were exposed to CyB or CyB-free clean water for four days, and the morphological endpoint was detected as described above (Duan et al., 2021). The daily mortality, 24h voluntary movement, heartbeat, body length, hatching rate and other developmental status were evaluated.

### 2.4 Gonadal histological examination

Histological specimens of F0 gonad were prepared according to previous methods (Qian et al., 2020) and histological examination was under the olympus microscope (Olympus, Japan).

### 2.5 E2, T, 11-KT and VTG concentrations measurement

Enzyme-linked immunosorbent assays (ELISA) kit (Yanjin Biological Co., Ltd., Shanghai, China) was used for determination. Briefly, blood samples were centrifuged (4°C, 3000 rpm) to obtain plasma, which was diluted with ELISA buffer to determine sex steroid hormones. See supplementary information for specific operation methods.

### 2.6 Gene Expression analysis

Total mRNA was extracted from brain, liver and gonad tissues by TRIzol reagent protocol (Tiangen Biotech, China). The first strand cDNA was synthesized by FasQuant RTase kit (Tiangen Biotechnology, China), the gene expression was measured by SYBR Green PCR Master Mix kit, and β-actin was used as the housekeeping gene. See supplementary information (Table S1) for details of primer sequence and RT-qPCR.

### 2.7 Statistical Analysis

SPSS 17.0 software (SPSS, USA) and GraphPad Prism 6.0 (GraphPad Software Inc.) were used for data analysis and image drawing. Significant differences between the control and CyB-treated groups were analyzed by one-way ANOVA followed by Dunnett’s test. P value < 0.05 was considered as statistically significantly different.

## 3. Results

### 3.1 Development of the F0 Zebrafish

The effect on the growth and development of F0 adult zebrafish after 150 days of CyB exposure is shown in Figure 1. K-factor, BSI, HSI and GSI of zebrafish were not significantly affected.

**Fig. 1.**
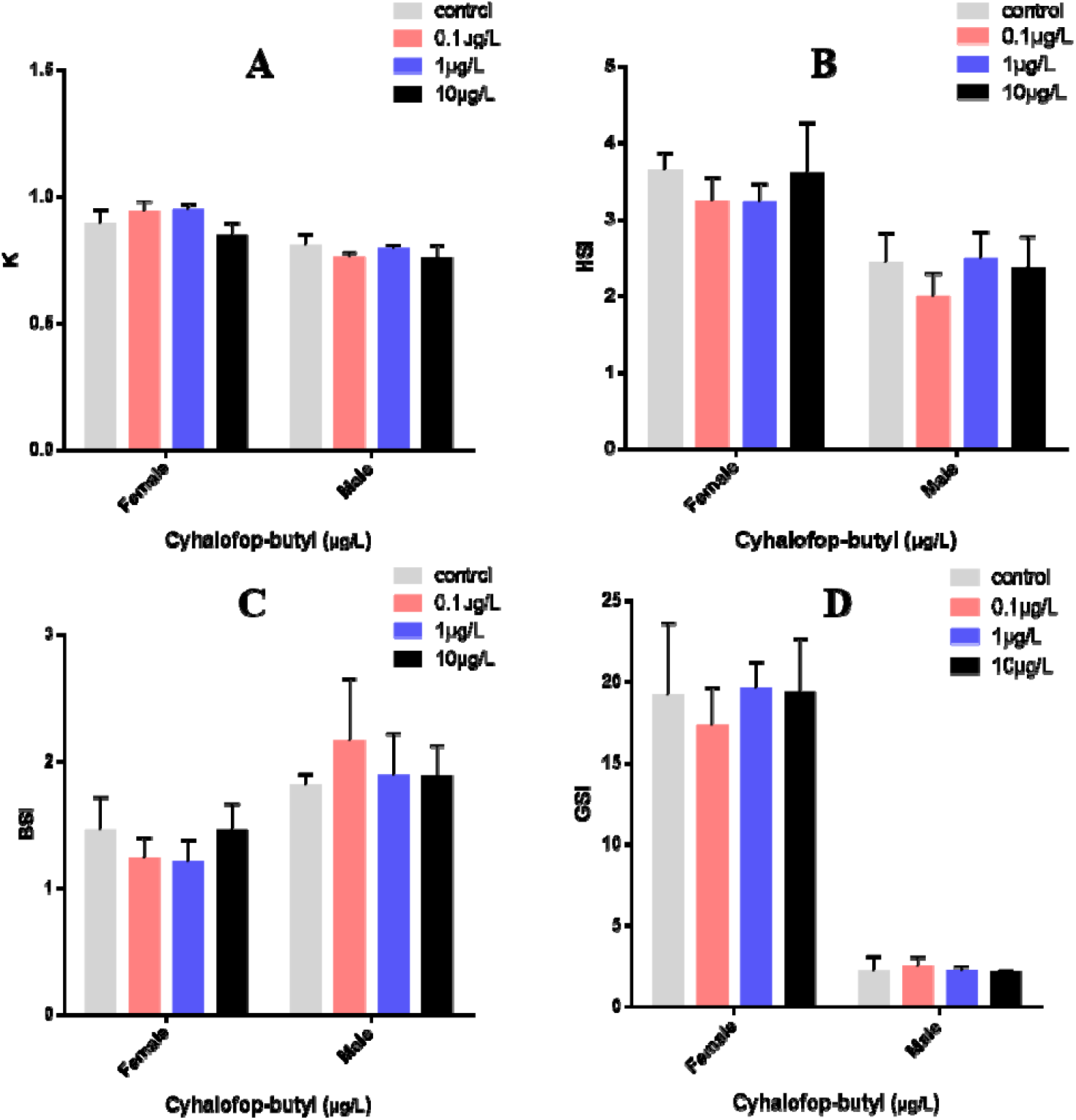
Effects of CyB on the growth and development of F0 adult zebrafish after 150 days. A: K = weight(g)/length3(cm) × 100. B: HSI = liver weight × 100/total. C: BSI = brain weight × 100/total weight. weight. D: GSI = gonad weight × 100/total weight. The asterisk indicates significant difference between control group and exposure groups (P < 0.05).

### 3.2 Measurement of F0 fertility and F1growth and development

After exposure to CyB, the fecundity of F0 zebrafish decreased significantly in a concentration-dependent manner (Figure 2A), and the unfertilized parent zebrafish exposed to 1 and 10 ug/L CyB also increased (Figure 2B). In addition, the F1 offspring of zebrafish treated with CyB developed abnormally, and the abnormal development of F1 generation was relieved when exposed to clear water without CyB (Figure 3). The embryo mortality of F1 zebrafish treated with CyB increased significantly with the increase of concentration (Figure 3A). Hatching rate did not change significantly (Figure 3B). Autonomy and body length decreased with the increase of CyB concentration, and the body length decreased significantly in the highest concentration group (10 ug/LCyB) where CyB continued to be exposed (Figure 3C-D). In the treatment groups of 1 and 10 ug/L CyB, the heart rate of Fl embryos exposed to CyB decreased significantly at 48, 72 and 96 hpf, while the heart rate decreased scarely when exposed to clean water without CyB, of which 72 hpf was the most effective (Figure 3E-F).

**Fig. 2.**
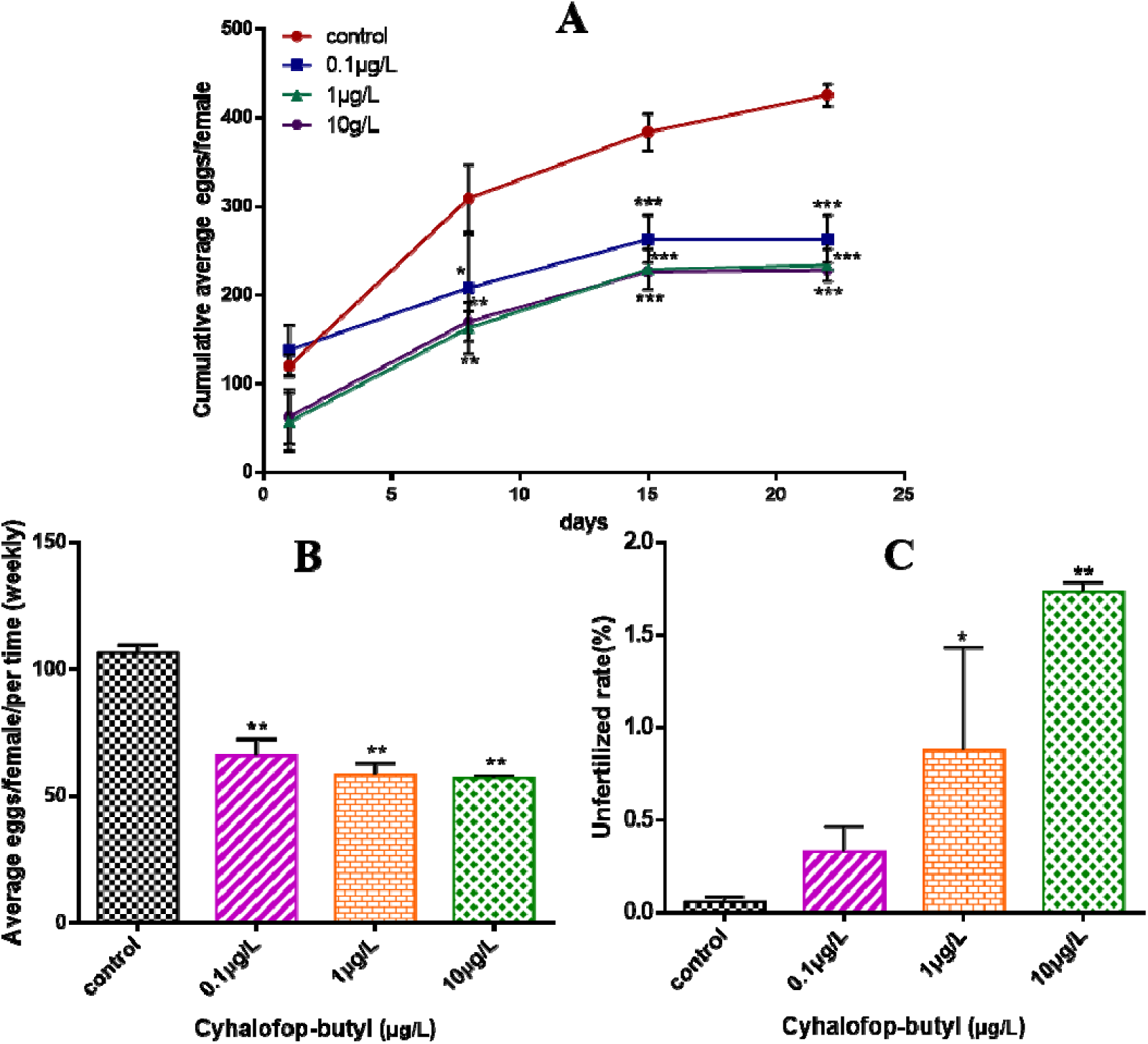
Effects of CyB on fertility of F0 zebrafish. A: Cumulative average number of eggs per female. B: Average number of spawning eggs per female and per time(weekly). (D) Unfertilization rate during the 150d of exposure. Unfertilization rate was calculated as cumulative number of unfertilized eggs × 100/total number of spawning eggs. Values are shown as the mean ± SD of three replicates per treatment (*p < 0.05; **p < 0.01).

**Fig. 3.**
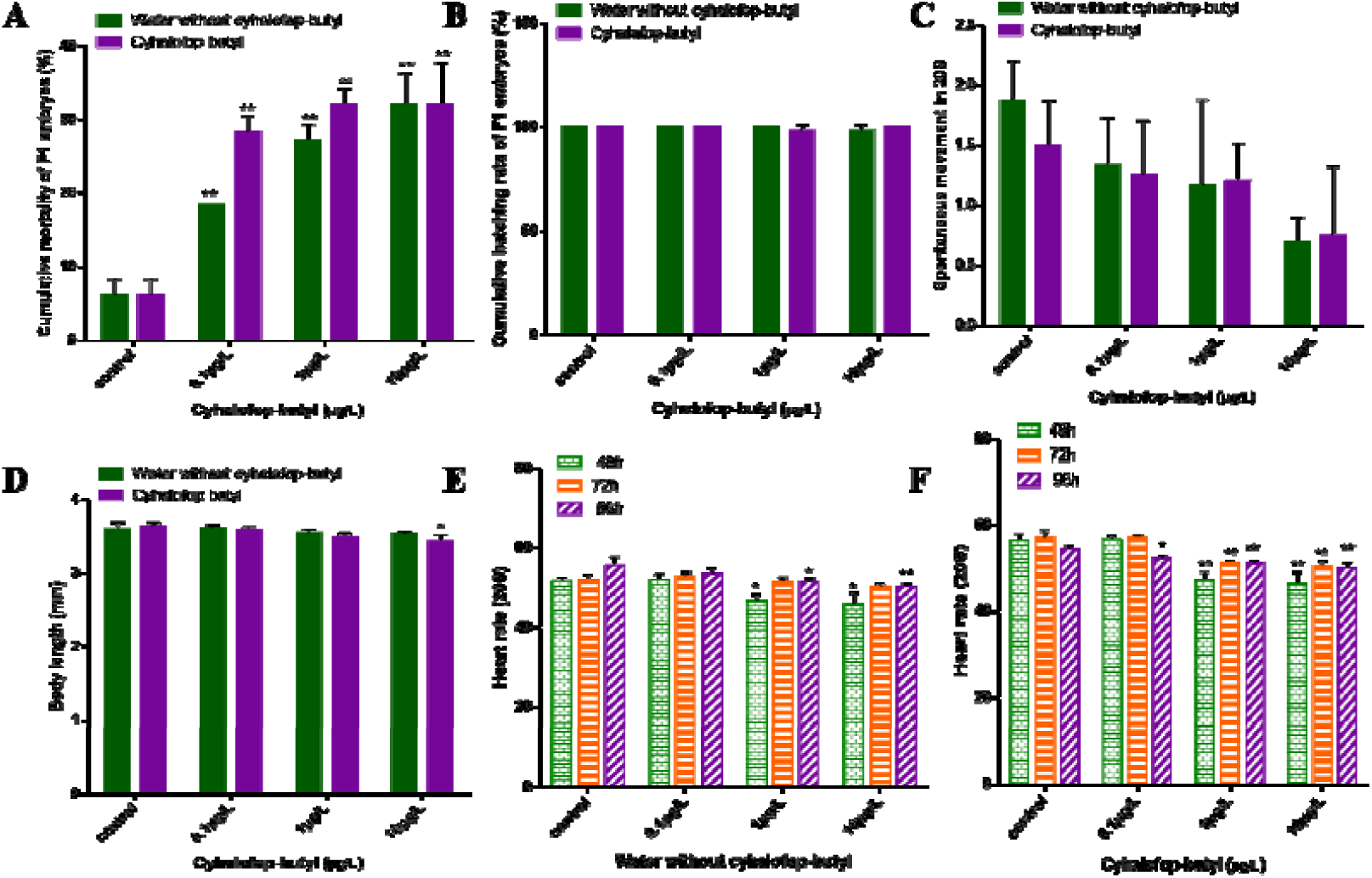
Effects of continuous exposure to CyB and water without CyB on embryo development of F1 zebrafish. (A) death rate at 96hpf, (B) hatching rate at 96 hpf, (C) spontaneous movement at 24 hpf, (D) body length at 96 hpf, (E-F) heartbeat at 48, 72 and 96 hpf. Asterisks marked the significant differences between control and CyB-treated groups (P < 0.05).

### 3.3 Gonadal histological examination

In female ovaries, there were no significant changes in perinuclear oocytes (PO), cortical alveolar oocytes (CO), early vitellogenic oocytes (EV) and late vitellogenic oocytes (LV) aall groups. In male testis, CyB treatmentdecreased the percentage of sperm (mature sperm cells) and spermatocytes, but there was no significant change (Figure 4F). The percentage of sperm cells (immature sperm cells) induced by 10 μg/L CyB increased significantly. A significant increase in the relative percentage of spermatogonia was observed in the 1 and 10 μg/L CyB groups (Figure 4G). The female gonads are mainly characterized by the separation of follicular wall and yolk or the detachment of outer membrane (Figure 4A-D), while male gonads are mainly characterized by widened stroma and decreased sperm number (Figure 4E).

**Fig. 4.**
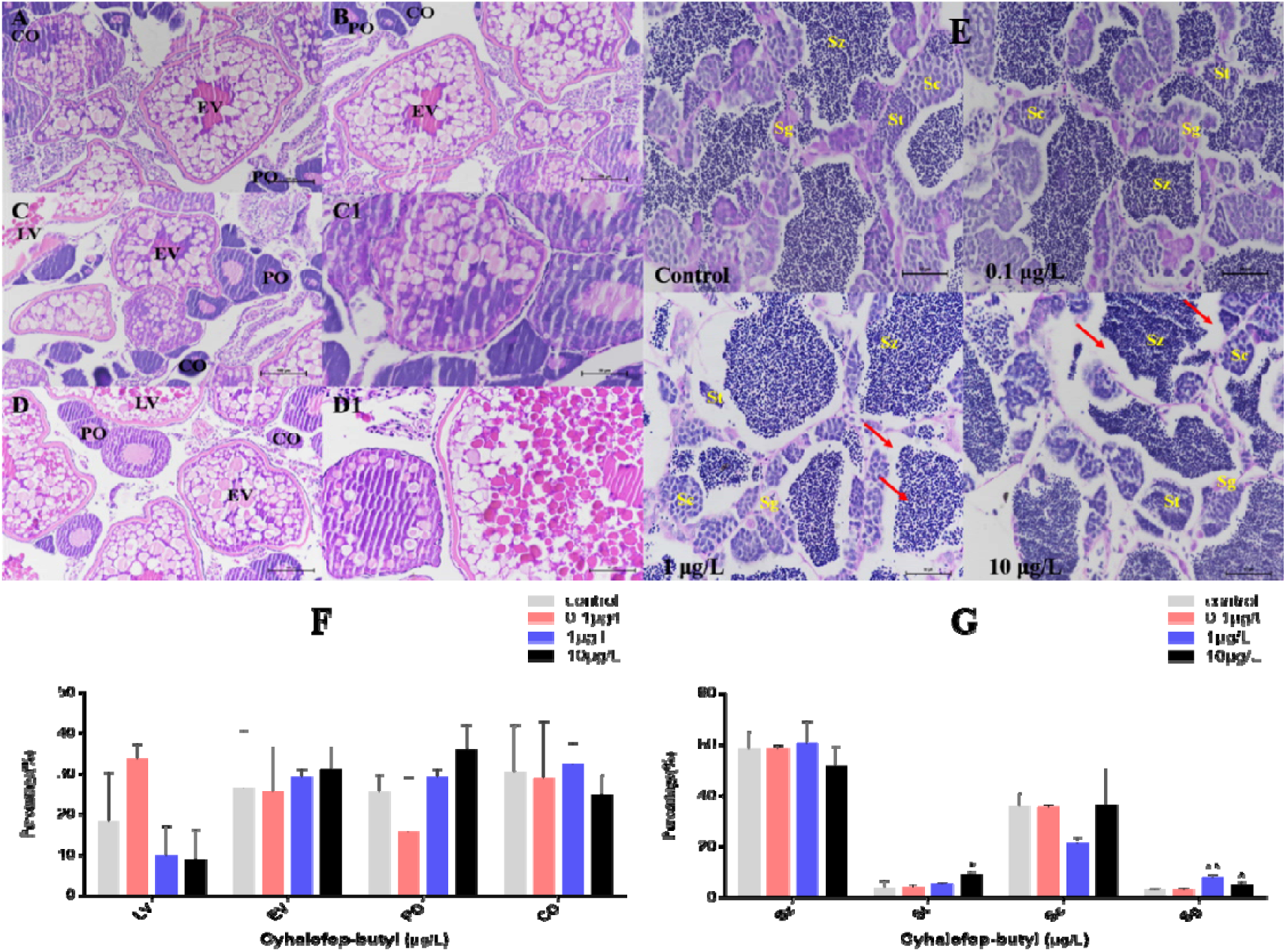
Histological observation of gonads of zebrafish exposed to CyB for 150 d. Females: (A) control; (B) 0.1 ug/L; (C/C1) 1 ug/L; (D/D1) 10 ug/L.The oocytes in the ovaries included perinucleolar oocytes (PO), cortical alveolar oocytes (CO), early vitellogenic oocytes (EV), and late vitellogenic oocytes (LV). (A/B/C/D, 200× magnification; C1/D1, 400× magnification). Males (E): control; 0.1 ug/L; 1 ug/L; 10 ug/L. The spermatocytes included spermatogonia (Sg), spermatocytes (Sc), spermatids (St), and spermatozoa (Sz), Red arrows indicate widened interstitial space. (400× magnification). Percentage (%) of different stages of oocytes in females (F) and spermatogenic cells in males (G). Results are presented as mean ± SD of three replicate samples (*p < 0.05; **p < 0.01).

### 3.4 Sex steroid hormone and VTG contents

In the plasma of F0 female zebrafish, the level of E2 decreased significantly, in all treatment groups, and that of male zebrafish only decreased significantly in 1 and 10 μg/L concentration groups (Figure. 5A). T level of female zebrafish significantly increased in all CyB exposed groups, while T level of male zebrafish significantly increased in the highest concentration group (10 μg/L) (Figure 5B). The 11-KT and VTG of zebrafish were significantly decreased in 1 and 10 μg/L concentration groups (Figure. 5C-D).

**Fig. 5.**
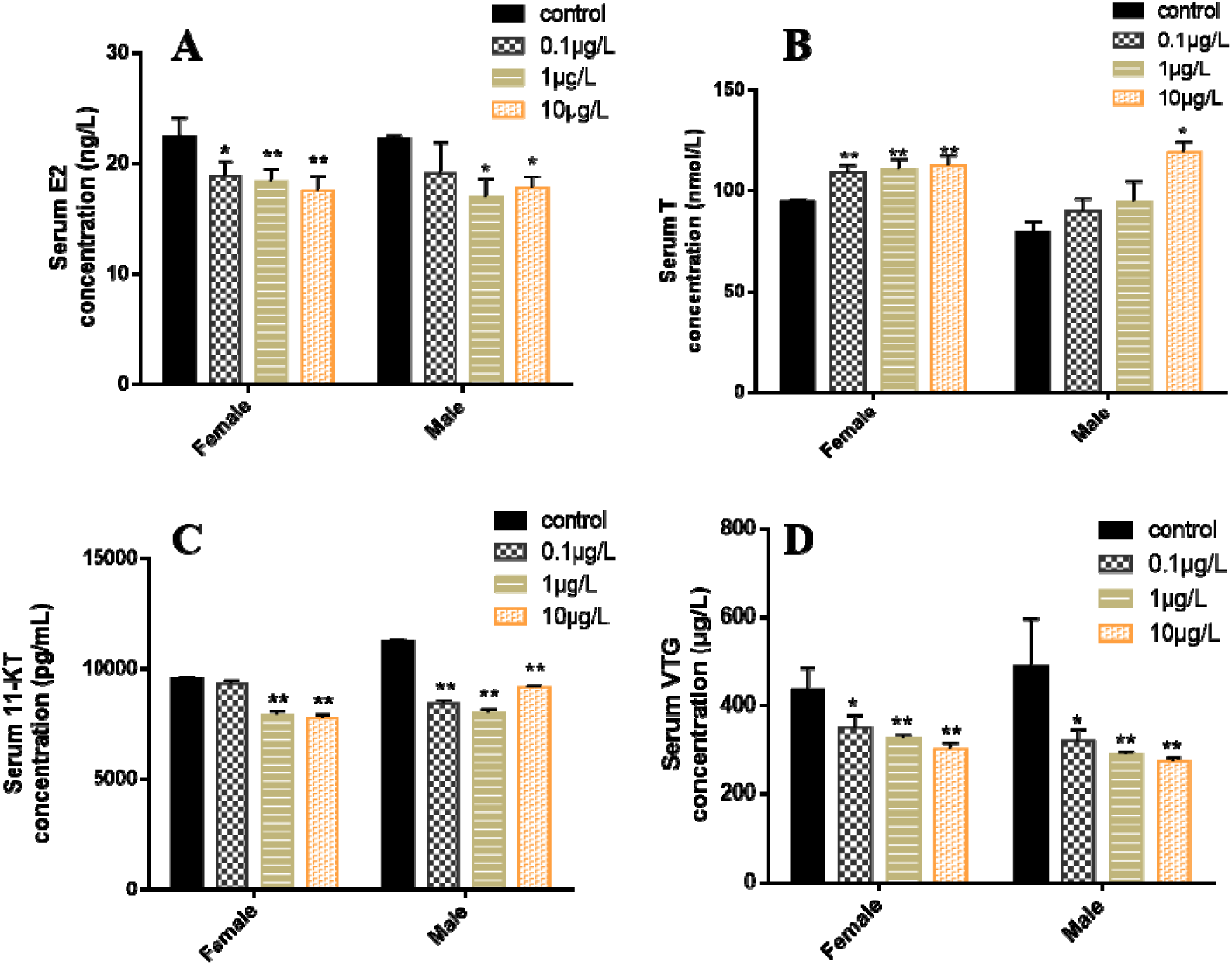
Effect of CyB exposure for 150 d on sex steroid hormones in F0 zebrafish. (A) 17β-estradiol (E2), (B) testosterone (T), (C) 11-ketotestosterone (11-KT) and (D) vitellogenin. Results are presented as mean ± SD of three replicate samples (*p < 0.05; **p < 0.01).

### 3.5 Gene expression alteration related to the HPGL axis

In the female zebrafish brains, the expression of *gnrhr2* and *lhb* has no significant chang after 0.1, 1 or 10 ug/L CyB exposure for 150 d compared with the control group, while the expressions of *gnrh2* and *gnrhr3* significantly increased in the 0.1 ug/L treatment group, *gnrh3, ar* and *esr2b* significantly down-regulated in the highest concentration treatment group (10 ug/L) (Fig. 6A). In F0 male brain, *gnrh3, ar, esr1*, and *esr2b* were increased but not significantly changed compared to controls, even in the highest concentration, while *gnrh2* was significantly down-regulated in all treatment group and *lhb* significantly increased at 1 and 10 ug/L groups. The expression of *cyp19b* and *fshb* was significantly up-regulated in the 10 ug/L CyB group (Fig. 6B). Neither *vtg1* nor *vtg2* levels were significantly changed in the livers of F0 females in the CyB-treated group compared to the control group (Fig. 6C). In F0 male livers, 10 ug/L CyB significantly down-regulated the expression levels of *vtg1* and *vtg2* (Fig. 6D). In F0 female gonads, compared with the control group, *hsd3b, fshr* and *lhr* did not change significantly, the expression of *cyp11a* was significantly up-regulated in the 1 and 10ug/L CyB groups, and the expression of *cyp19a* and *hsd17b* in the highest concentration treatment group (10ug/L) was significantly upregulated (Fig. 6E). In F0 male gonads, *cyp11a* was significantly up-regulated in 1 and 10 ug/L treatment groups, *cyp19a* and *fshr* were significantly up-regulated in 0.1 ug/L CyB group, and *cyp19a* and *hsd17b* were significantly down-regulated in 10 ug/L CyB group, the expressions of *hsd17b* and hsd3b were significantly up-regulated in 0.1 and 1 ug/L CyB groups (Fig. 6F).

**Fig. 6.**
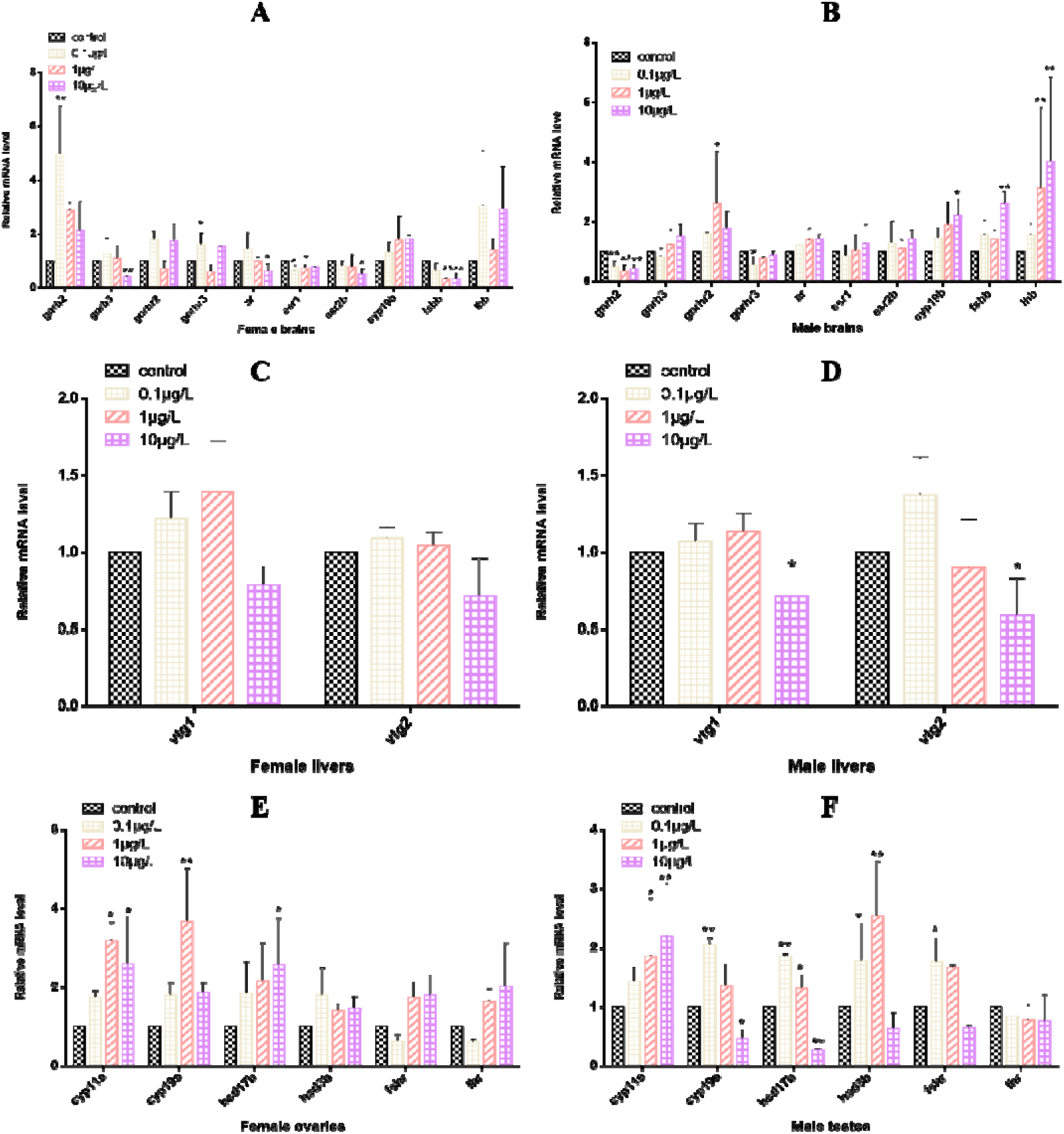
Transcri ption of HPGL axis related genes in F0 zebrafish after 150 days of exposure to CyB. female: (A) brain, (C) liver and (E) gonad; Male: (B) brain, (D) liver and (F) gonad. Results are presented as mean ± SD of three replicate samples (*p < 0.05; **p < 0.01).

## 4. Discussion

The reproductive capacity of aquatic organisms is of great significance to the maintenance of population density and the stability of aquatic ecosystem (Arcand-Hoy and Benson, 1998; Nassar and Fahmy, 2015). However, many environmental pollutants, especially pesticides (such as glyphosate, dimethylbenzene and pretilachlor), have been reported to have significant inhibitory effects on the reproductive capacity of fish in rivers and oceans (Gupta and Verma, 2020; Soni and Verma, 2020; Uren Webster et al., 2014). Similarly, in this study, it was found that cyhalofop-butyl had a significant inhibitory effect on the reproductive capacity of zebrafish. During the whole life cycle of zebrafish embryos exposed to CyB, the gonad index (GSI) of both male and female zebrafish showed a downward trend. The herbicide changes the content of sex hormones and vitellogenin to different degrees, and the relative expression of genes related to HPGL axis was disturbed, which affected the reproduction of the parent zebrafish, leading to the decline of reproductive capacity and fertilization capacity. In addition, harmful effects of parental zebrafish can be transferred to offspring, resulting in abnormal embryo development of F1 zebrafish.

The results showed that after exposure to CyB for 120d, the average accumulated egg production of zebrafish decreased significantly with the extension of exposure time under the environment-related concentration of CyB (0.01μg/L), and severe reproductive inhibition was observed in the treatment groups with high concentrations of CyB (1 and 10 μg /L). The relative oviposition of zebrafish decreased by 45% and 46% during the whole exposure period, respectively. Although GSI is usually a quantitative indicator of sexual maturity and ovarian development of vertebrates (Pyle et al., 2005; Van den Belt et al., 2002), the value of GSI is often affected by other factors, such as oviposition cycle, the determination of GSI may not be as sensitive as gonad histology analysis in the evaluation of reproductive toxicity. In the study, compared with the control group, the fecundity in 10 μg /L CyB exposed group decreased significantly, and the percentage of LV decreased by 9%, while the GSI value did not change significantly. A similar change was observed after 21 days of exposure to zebrafish with 1mg/L boscalid. The female fish’s fertility decreased, and the oocyte stage distribution in the ovary was abnormal, but the GSI value did not change (Qian et al., 2020). At the same time, CyB exposure also significantly affected the reproductive system of male zebrafish. 1 μg/L and 10 μg/L CyB groups resulted in the inhibition of spermatogenesis in the testis of male fish. Inhibition of parental spermatogenesis may affect the fertilization rate of F1 generation. Similarly, the study found that the chronic exposure of azoxystrobin 200 μg/L also lead to inhibited spermatogenesis of zebrafish males and decreased fertilization rate of F1 generation (Cao et al., 2019). In addition, CyB exposure also caused tissue damage of zebrafish gonad, characterized by separation of female gonad follicle wall from yolk or shedding of outer membrane, widening of male gonadal stroma and decrease of sperm number.

In this study, E2 content in F0 zebrafish exposed to CyB decreased significantly, while T content increased, which indicated that the steady state of sex steroid hormone was destroyed. In teleost, the content and balance of E2 and T sex steroid hormones are considered to play an important role in sex differentiation and reproduction (Cao et al., 2016b). In addition, VTG is the energy and nutrients needed for the growth and development of newborn fish embryos and young fish (Navas and Segner, 2006). In fish, E2 can induce the production of VTG, and regulate the synthesis of VTG and its gene expression in liver (Chen and Chan, 2016; Flouriot et al., 1996). Therefore, the decrease of VTG content in male and female zebrafish may be induced by the significant decrease of E2 content. The decrease of E2 content in female and male fish may reduce vitellogenesis in ovary (Sundararaj et al., 1982), at the same time, this is consistent with the decrease of the expression of *vtg1* and *vtg2* in the liver of male and female fish. Therefore, the collective results show that CyB will disturb the hormone balance and affect the reproduction of zebrafish.

HPGL axis regulates the physiological process of fish gametogenesis, and the content of sex hormones is associated with the changes in sex hormone synthesis related genes regulated by HPGL axis (Phumyu et al., 2012; Weltzien et al., 2004). GnRH is a biosynthetic gonadotropin (GnHs) in hypothalamus, which is regulated by HPGL axis (Ji et al., 2013a; Teng et al., 2020). The increased expression of *gnrh2, gnrhr2* and *gnrhr3* in females and the up-regulation of *gnrhr3* and *gnrhr2* in males are consistent with the decrease of E2 production, which indicates that CyB can directly regulate the content of GnRH, thus affecting the secretion of GnHs. In the process of regulation, pituitary gland synthesizes and secretes key hormones on HPGL axis, such as follicle-stimulating hormone (FSH) and luteinizing hormone (LH), which promote ovarian development and differentiation, regulate gamete formation and steroid hormone synthesis (Hauser, 2014; Schulz et al., 2001). FSH is a glycoprotein that can promote E2 synthesis, gonadal hormone secretion and puberty spermatogenesis (Kwok et al., 2005), LH stimulates the synthesis of androgens and progesterone secretion (Anway, 2010; Qian et al., 2020; Schulz et al., 2010). Therefore, the decrease of *fsh* in zebrafish brain may inhibit the synthesis of E2 in female fish, resulting in the change of LV in ovary and the subsequent decrease of fecundity. The increase of *fshb* and *lhb* expression in male zebrafish may affect gonad development and change the percentage of St and Sg. In addition, the biosynthesis process of steroid hormones is directly related to steroid synthase. Cholesterol is converted into testosterone by a series of enzymes (*cyp11a, hsd3b, cyp17* and *hsd17b* encode steroid synthase), and finally into estradiol by aromatase (encoded by *cyp19a*). Therefore, the expression changes of genes related to steroid-producing enzymes may interfere with the balance of sex hormones (Eidem et al., 2006; Trant et al., 2001). In addition, *hsd17b* catalyzes the conversion of androstenedione to T, which then converts to 11-KT (Mindnich et al., 2005). The up-regulation of *hsd17b* in female ovary and the up-regulation and down-regulation of *hsd17b* in male testis showed that CyB interfered the steroid pathway and damaged the biosynthesis of sex hormones, thus increasing the level of T and decreasing the level of 11-KT, and destroying the reproductive system of male zebrafish. Aromatase (CYP19) is a ccrucial enzyme that catalyzes the conversion of androgen to estrogen in fish. It regulates sex differentiation and reproductive behavior of most teleost fish by influencing E2 synthesis. *cyp19a* is mainly expressed in gonad, and *cyp19b* is mainly expressed in brain (Cheshenko et al., 2007; Cheshenko et al., 2008). In this study, *cyp19b* was expressed more in male brain, indicating that the transformation from E2 to T was increased, resulting in the significant decrease of E2 and the significant increase of T in plasma. Therefore, we speculate that the decrease of *cyp19a* expression in gonad of male fish prevents testosterone from being converted into estradiol, which leads to the decrease of E2 synthesis in plasma and the increase of testosterone content. Previous studies have shown similar results, the exposure of azoxystrobin leads to the decrease of *cyp19a* expression in female zebrafish ovaries and the decrease of estradiol content in female zebrafish (Cao et al., 2016b). Tebuconazole suppressed the expression of *cyp19a* in HPGL axis of zebrafis, and decreased the content of estradiol in female fish (Li et al., 2019).

In addition, the harmful effects of the parent zebrafish can be transferred to the offspring, resulting in a decrease in the survival rate, heart rate and body length of F1 zebrafish embryos, even though F1 embryos have been relieved in clean water. Since fish offspring and their parents may continue to live in the same water environment in nature (Dong et al., 2018), when parents are exposed to pollutants, the offspring may be affected not only by direct exposure, but also by parental exposure (Aluru et al., 2010; Chen et al., 2017; Galus et al., 2014; Hurem et al., 2017; Liu et al., 2014).

## 5. Conclusion

In this study, life cycle exposure to CyB adversely effected the reproduction of zebrafish adults. The change of mRNA expression of HPGL axis-related markers may be the underlying molecular mechanism of zebrafish reproductive damage. In addition, the exposure of F0 zebrafish to CyB concentration related to the environment affected the survival rate, heart rate and body length of F1 zebrafish after continuous exposure and fresh water recovery. These data provide a new understanding of the effects of long-term exposure of zebrafish to environmental-related concentrations of CyB on reproduction.

## Acknowledgments

This work was supported by the Advanced Talents Incubation Program of Hebei University (No. 050001-521000081469).

## Graphic Abstract

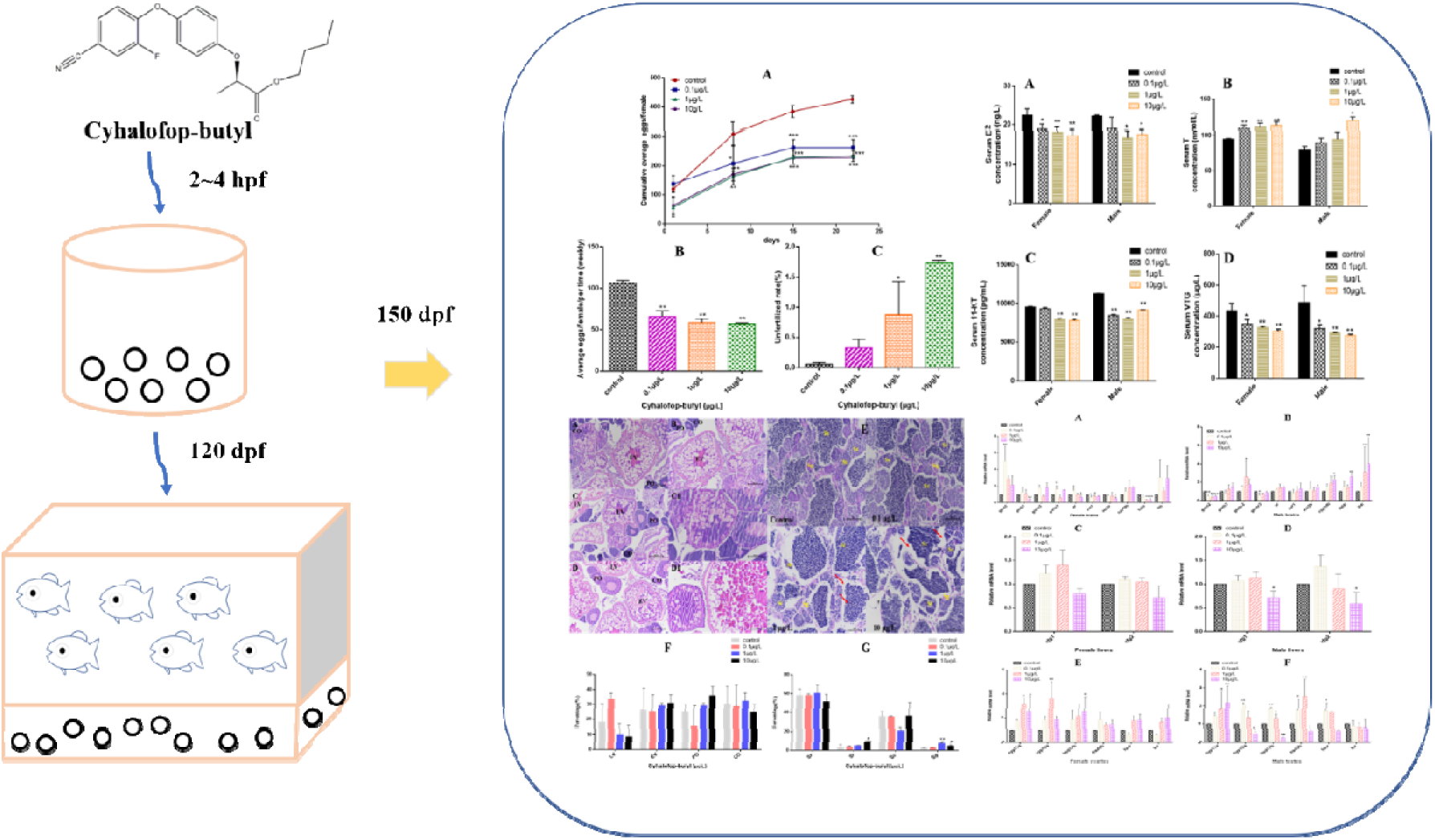

