## Supplementary Information for "Life cycle exposure to cyhalofop-butyl induced reproductive toxicity toward zebrafish"

No. 180 Wusi East Road, Baoding City, Hebei Province

### Appendix S1. The method of sex steroid hormone and VTG concentrations measurement

The serum samples used to measure E2, T, 11-KT and VTG were centrifuged (4 °C, 3000 rpm, 10 min), and 10 μL of the supernatant was pipetted into a 96-well microplate. Enzyme-linked immunosorbent assay (ELISA) kit (Yanjin Biological Co., Ltd., Shanghai, China) was used to dilute the supernatant twice for the following measurement. The procedure is as follows:

1) add 10 μL of diluted serum supernatant and 40 μL of sample diluent into antibody-coated microtiter plates.

2) after a series of incubation, washing plates, enzyme catalysis, incubation, washing plates, color rendering and termination, the absorbance was measured at 450 nm using a microplate reader.

3) calculate the actual concentrations of E2, T, 11-KT and VTG of serum sample according to the standard curve.

Table S1. The sequence of primers related to the HPGL axis.

| Primer 名称 | 序列(5' to 3') |
|---|---|
| β-actin F | TGGACTCTGGTGATGGTGTGAC |
| β-actinR | GAGGAAGAAGAGGCAGCGGTTC |
| gnrh2 F | GGTCTCACGGCTGGTATCCT |
| gnrh2 R | TGCCTCGCAGAGCTTCACT |
| gnrh3 F | TGGTCCAGTTGTTGCTGTTAGTT |
| gnrh3 R | CCTGAATGTTGCCTCCATTTC |
| gnrhr2 F | ACAGCGTGAGCAAAACATTG |
| gnrhr2 R | TGAGCACAAACTCAGCATCC |
| gnrhr3 F | AACAGACATGATCCCGAAGG |
| gnrhr3 R | AGGTTCCCGAACACAAACAG |
| esr1 F | CCCACAGGACAAGAGGAAGA |
| esr1 R | CCTGGTCATGCAGAGACAGA |
| esr2b F | CAACAGGGAGGAAGGGAA |
| esr2b R | TTAGCAGATGAGCGAGCC |
| ar F | ACATTCTGGAGGCCATTGAG |
| ar R | ACGTGCAAGTTACGGAAACC |
| vtg1 F | CTGCGTGAAGTTGTCATGCT |
| vtg1 R | GACCAGCATTGCCCATAACT |
| vtg2 F | TACTTTGGGCACTGATGCAA |
| vtg2 R | AGACTTCGTGAAGCCCAAGA |
| cyp11a F | AATGGGAAGTATCCTGGTG |
| cyp11a R | CTGTAGGTCTGGCTGTCG |
| cyp19a F | GCTGACGGATGCTCAAGGA |
| cyp19a R | AAACGTCCACCACGATGCA |
| cyp19b F | CAGTCGTTACTTCCAGCCATTC |
| cyp19b R | CCGCTGTTTCTCCGTTGC |
| hsd17b F | ACATTCACGGCTGAGGAGTTT |
| hsd17b R | ATGCTGCCATACGTTTGGTC |
| hsd3b F | GCAACTCTGGTTTTCCACACTG |
| hsd3b R | CAGCAGGAGCCGTGTAGCTT |
| fshr F | CGTCTCTTTTGTGCACTGGA |
| fshr R | GTGGCAATTCCACACTTCCT |
| lhr F | CCTGGTCGTCCTGCTGGTT |
| lhr R | AAGGCTAGATGGCACATTAGAAATC |
| fshb F | GCAGGACTATGCTGGACAATG |
| fshb R | CCACGGGGTACACGAAGACT |
| lhb F | GGCTGGAAATGGTGTCTTCTT |
| lhb R | GGAAAACGGGCTCTTGTAAAC |
